## Supplementary Information for "X-chromosome upregulation operates on a gene-by-gene basis at RNA and protein levels"

**Supplementary Figure 1-9**

**Supplementary Table 1-6**

### Supplementary Figure 1

A

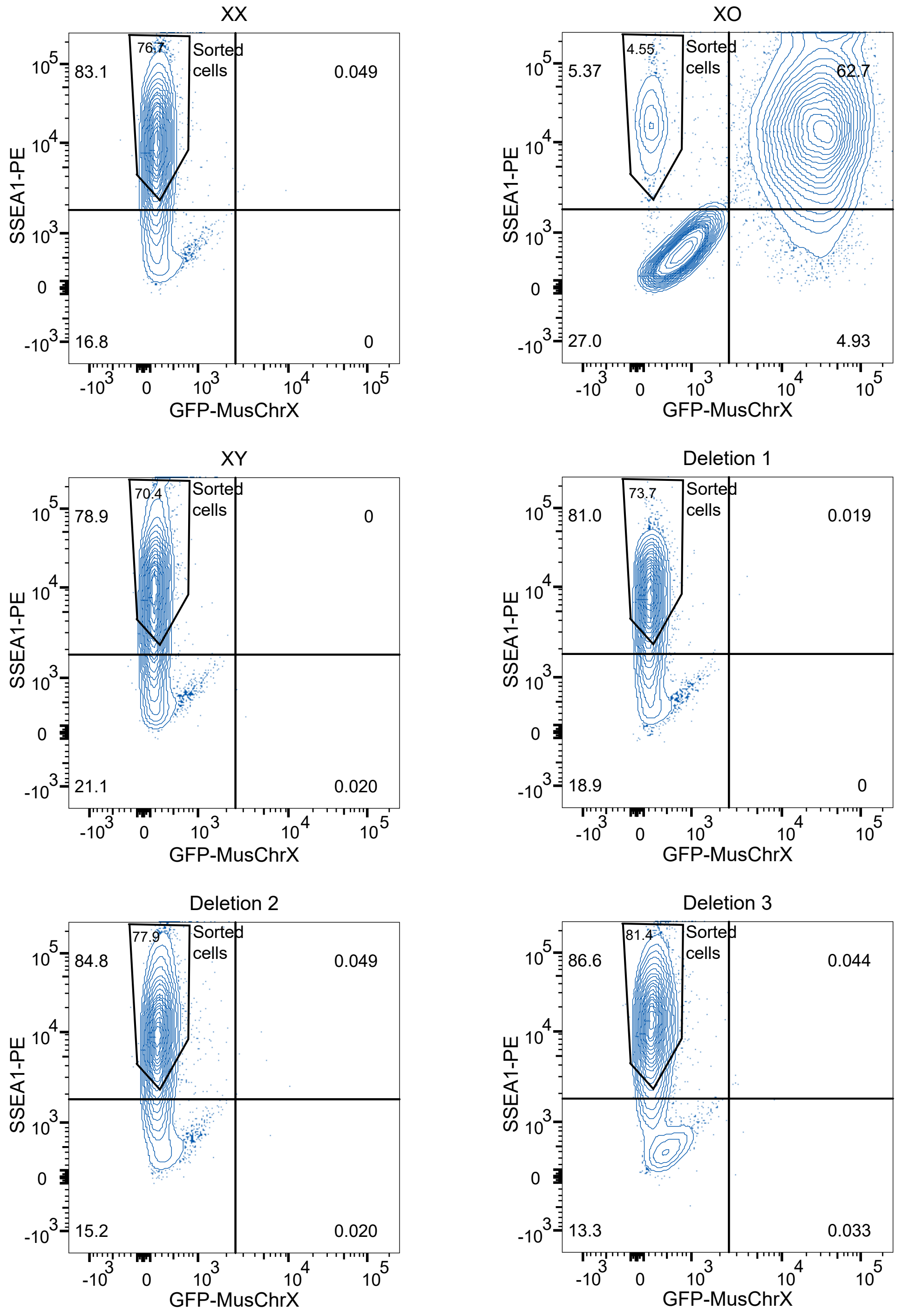


B

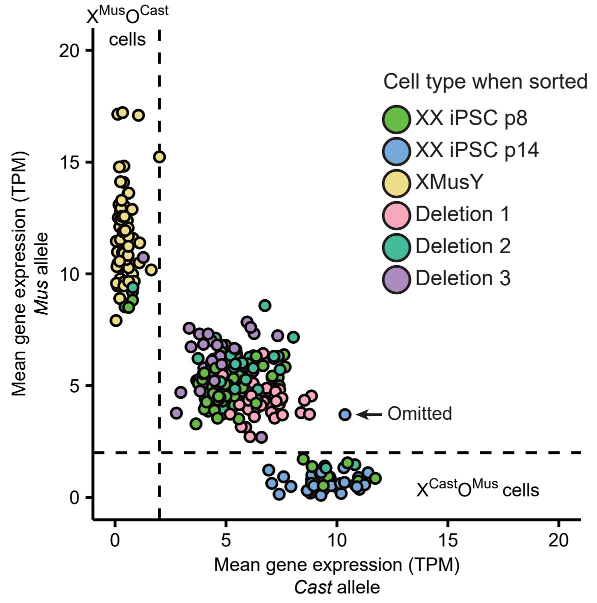

**Supplementary Fig. 1. FACS sorting and classifying single cells after sequencing.**
**a.** Flow cytometry contour plots of passage 8 iPSCs (XX), passage 14 iPSCs (XO), XY mESCs, and cells with Deletion 1, 2 and 3. Cells were sorted for positive SSEA1 signal and no transgenic GFP expression.
**b.** Scatter plot of average X-linked gene expression from each allele for each cell. Cells are colored by identity when sorting. Each dot represents a cell. Dashed lines indicate thresholds for defining a cell as XO: Cast expression > 2 (vertical line) and Mus expression > 2 (horizontal line). One XX p14 cell was omitted from the dataset due to ambiguous levels of expression from both X chromosomes.

### Supplementary Figure 2

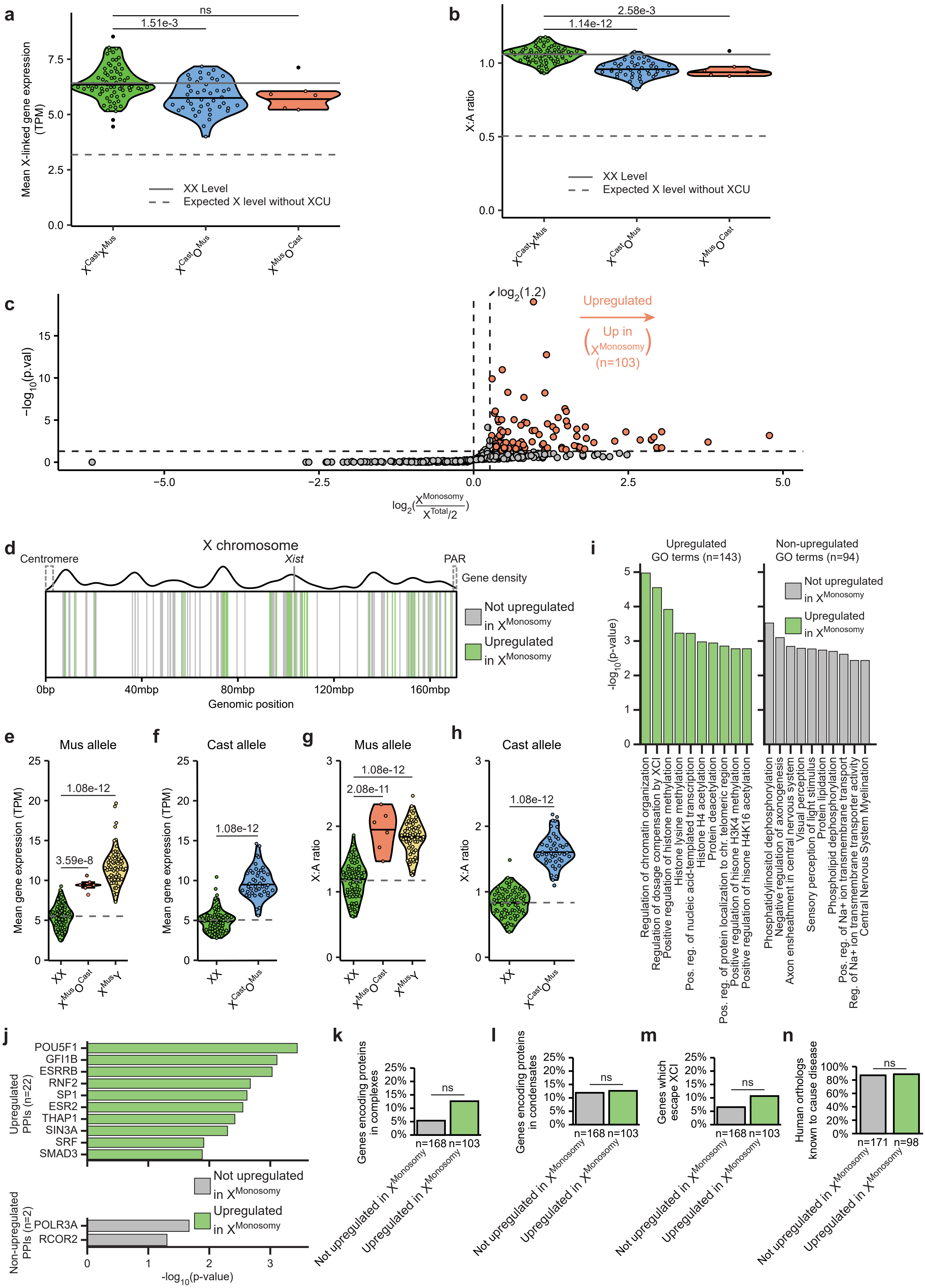

**Supplementary Fig. 2. Identification of upregulated genes in cell lines with total X-chromosome monosomy.**

**a.** Violin plots of mean X-linked gene expression per cell for $X^{Cast}X^{Mus}$, $X^{Cast}O^{Mus}$, and $X^{Mus}O^{Cast}$ cells. The solid line indicates the mean value of $X^{Cast}X^{Mus}$ cells, and the dashed line indicates half of the mean value of $X^{Cast}X^{Mus}$ cells, serving as a reference line for expected expression without upregulation. Each dot represents a cell. P-values were calculated via the Tukey HSD test.

**b.** Violin plots of X-to-Autosome (X:A) ratio per cell for $X^{Cast}X^{Mus}$, $X^{Cast}O^{Mus}$, and $X^{Mus}O^{Cast}$ cells. The solid line indicates the mean value of $X^{Cast}X^{Mus}$ cells, and the dashed line indicates half of the mean value of $X^{Cast}X^{Mus}$ cells, serving as a reference line for expected expression without upregulation. Each dot represents a cell. P-values were calculated via the Tukey HSD test.

**c.** Volcano plot of X-linked genes in X-monosomic cells ($X^{Cast}O^{Mus}$, $X^{Mus}O^{Cast}$, $X^{Mus}Y$ together) and $X^{Cast}X^{Mus}$ cells. Each dot represents a gene. P-values are calculated via one-sided Student's t-test with alternative hypothesis of $X^{monosomy} > X^{Cast}X^{Mus}*0.5$. Horizontal dashed lines represent *P-value* = 0.05; vertical dashed lines represent fold change = 1.2. $X^{Cast}X^{Mus}$ expression values are copy number-corrected (50% of total allelic expression).

**d.** Schematic depiction showing the locations of all upregulated (green) and non-upregulated (gray) genes, as well as the location of the centromere, *Xist* locus, and pseudo-autosomal region (PAR), and the gene density across the X chromosome.

**e, f.** Violin plots of the mean normalized gene expression from the *Mus* (**e**) and Cast (**f**) allele of upregulated genes in $X^{Cast}X^{Mus}$, $X^{Cast}O^{Mus}$,$X^{Mus}O^{Cast}$, and $X^{Mus}Y$ cells. Dashed line indicates the mean value of XX cells, serving as a reference line for expected expression without upregulation. Each dot represents a cell. P-values were calculated via the Tukey HSD test.

**g, h.** Violin plots of the mean X:A ratio of upregulated genes of the *Mus* (**g**) and Cast (**h**) allele in $X^{Cast}X^{Mus}$, $X^{Cast}O^{Mus}$,$X^{Mus}O^{Cast}$, and $X^{Mus}Y$ cells. The dashed line indicates the mean value of XX cells, serving as a reference line for expected expression without upregulation. Each dot represents a cell. P-values were calculated using the Tukey HSD test.

**i.** GO enrichment analysis on upregulated (green, left) and non-upregulated (gray, right) gene lists. The p-values of the top 10 GO terms in each gene list are shown separately. Total number of upregulated terms per gene list is displayed on the top. Pos. reg. = positive regulation.

**j.** Enrichment of transcription factor protein-protein interactions (PPI) in the upregulated (green, top) and non-upregulated (gray, bottom) gene lists. The p-values of the top 10 significant transcription factors in each gene list are shown separately. Total number of upregulated transcription factors per gene list is displayed on the left.

**k-n.** Bar plots showing the percentage of genes which code for complex-forming proteins (**k**), proteins which form condensates (**l**), genes which escape XCI (**m**) and genes with human orthologs known to cause disease (**n**) in the upregulated (green) and non-upregulated (gray) gene lists. P-values calculated via the Chi-square test (ns - not significant).

### Supplementary Figure 3

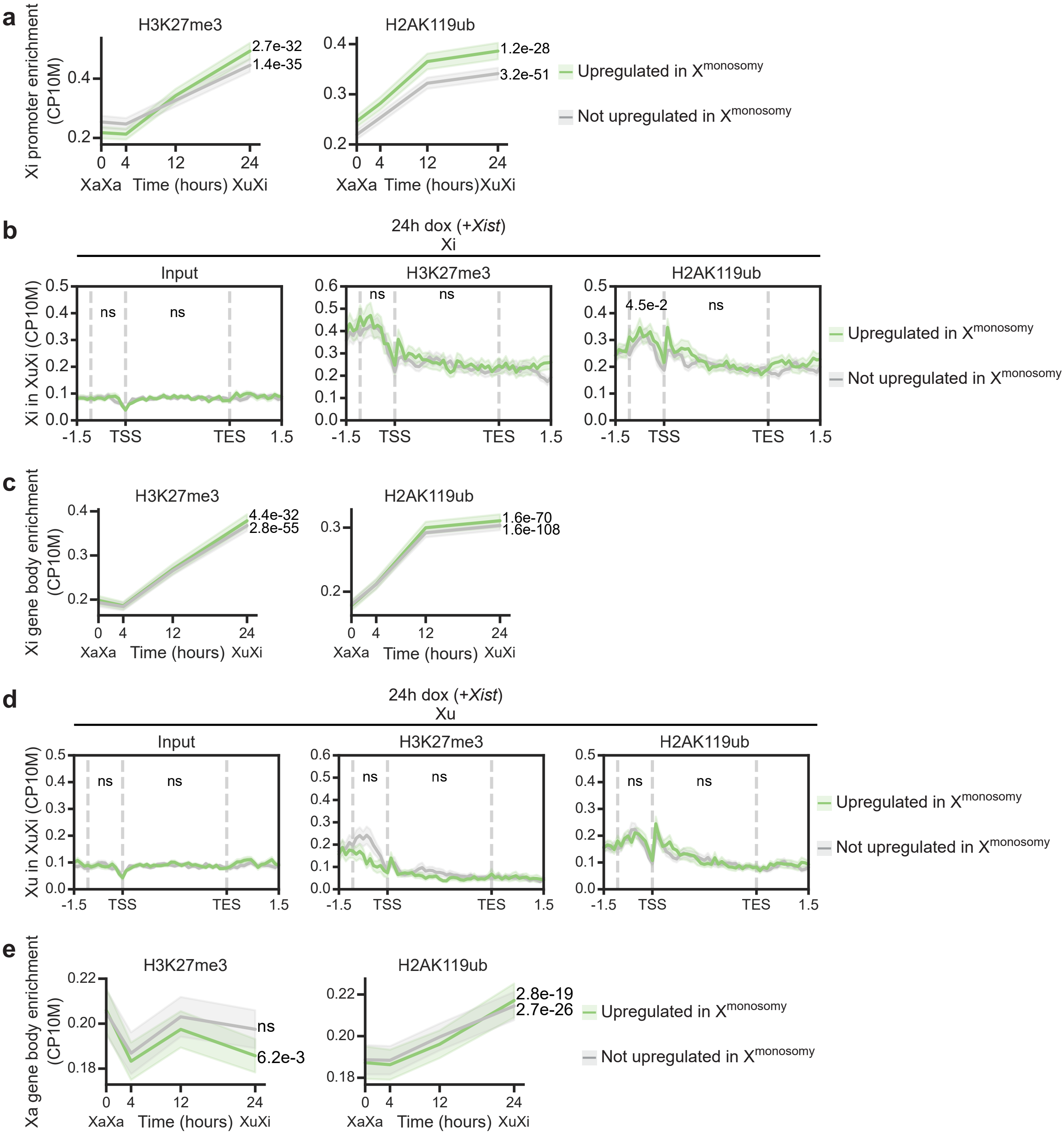

**Supplementary Fig. 3. Repressive hPTMs analysis.**

**a, c.** Line plots showing promoter (**a**) and gene body (**c**) enrichment of repressive histone modifications on the inactive allele during XCI. Enrichment was quantified by the mean counts per 10 million (CP10M) in the promoter region (1kb region upstream of the TSS, **a**) or gene body (TSS to TES, **c**). Mean enrichment is plotted for upregulated (green) and not upregulated (gray) genes separately, with shaded areas indicating the standard error. P-values comparing enrichment at 0 and 24 hours were calculated using two-sided paired t-tests, corrected for multiple testing with the Benjamini-Hochberg method, and shown to the right of each gene set.
**b, d.** Profile plots showing ChIP-seq enrichment at genomic regions of upregulated (green) and not upregulated genes (gray). The CP10M-normalized tracks of the inactive (**b**) or active upregulated (**d**) allele after 24 hours of dox are plotted, with shaded areas indicating the standard error. Promoter and gene body regions are indicated by dashed lines. P-values comparing enrichment between upregulated and not upregulated genes within these regions were calculated using two-sided Student's t-tests, corrected for multiple testing with the Benjamini-Hochberg method, and are shown above the corresponding region.
**e.** Same as in **c**, but showing gene body enrichment on the active upregulated allele.
Reanalysis of data from Zylicz et al.[57]

**Supplementary Figure 4**

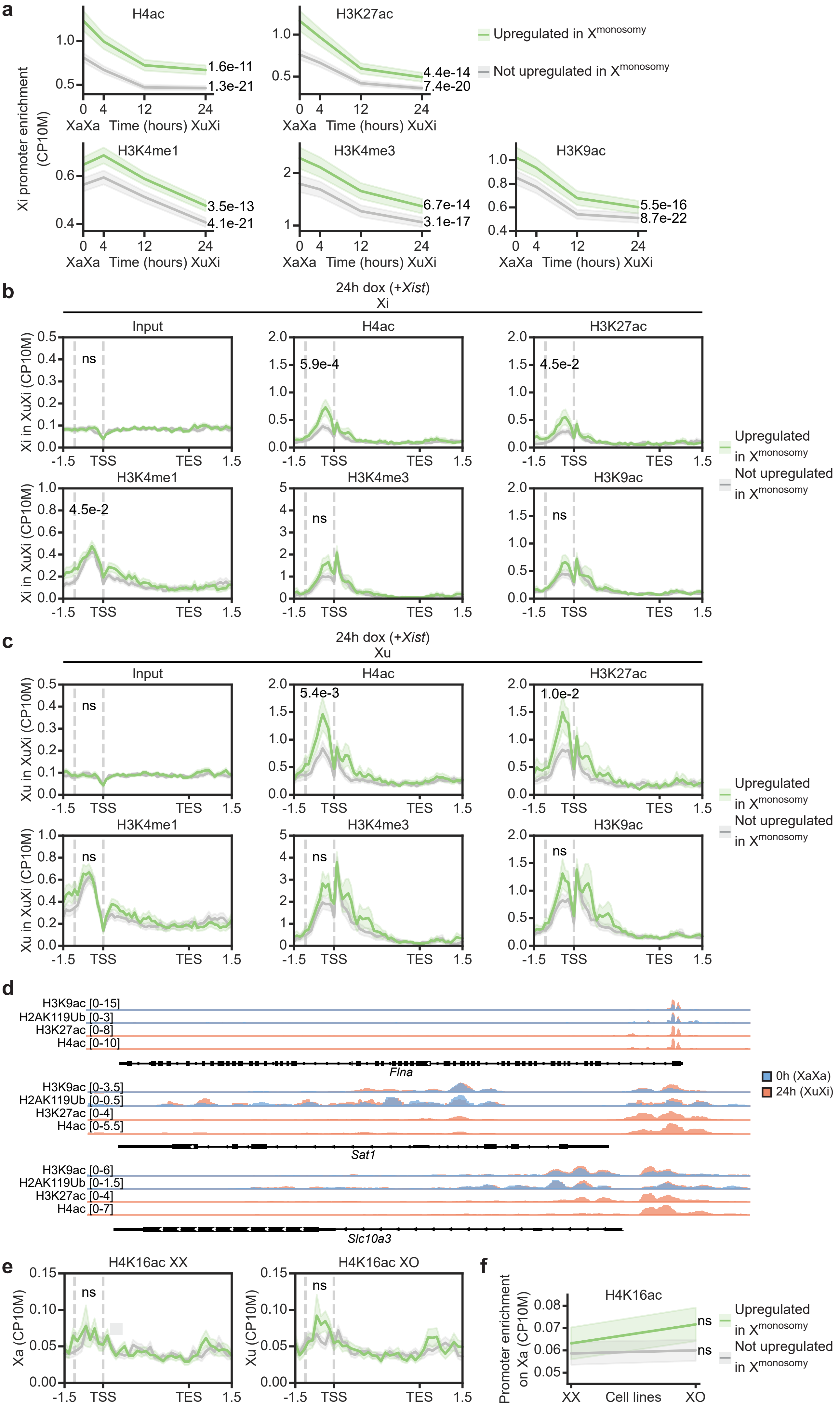

**Supplementary Fig. 4. Active hPTMs analysis.**

**a.** Line plots showing promoter enrichment of active histone modifications on the inactive allele during XCI. Promoter enrichment was quantified as the mean counts per 10 million (CP10M) in the promoter region (1kb region upstream of the TSS). Mean enrichment is plotted for the promoters of upregulated (green) and not upregulated (gray) genes separately, with shaded areas indicating the standard error. P-values comparing enrichment at 0 and 24 hours were calculated using two-sided paired t-tests, corrected for multiple testing with the Benjamini-Hochberg method, and shown to the right of each gene set.

**b-c**. Profile plots showing ChIP-seq enrichment at genomic regions of upregulated (green) and not upregulated genes (gray). The CP10M-normalized tracks of the inactive allele (**b**) or active upregulated allele (**c**) after 24 hours of dox are plotted, with shaded areas indicating the standard error. P-values comparing enrichment between upregulated and not upregulated genes within the promoter region were calculated using two-sided Student's t-tests, corrected for multiple testing with the Benjamini-Hochberg method, and are shown above the promoter region (indicated with dashed lines).

**d**. Genome browser views showing histone modification tracks at loci of upregulated genes *Flna*, *Sat1* and *Slc10a3*. Promoter regions display increased enrichment of H3K9ac and H2AK119Ub between 0 hours (blue) and 24 hours (orange), consistent with significant changes shown in **Figure 1h-i**. Additionally, promoter enrichment of H3K27ac and H4ac is observed at 24h, as described in **c**.

**e**. Same as in **c**, but for H4K16ac enrichment on the active allele.

**f**. Same as in **a**, but for H4K16ac promoter enrichment on the active allele.

### Supplementary Figure 5

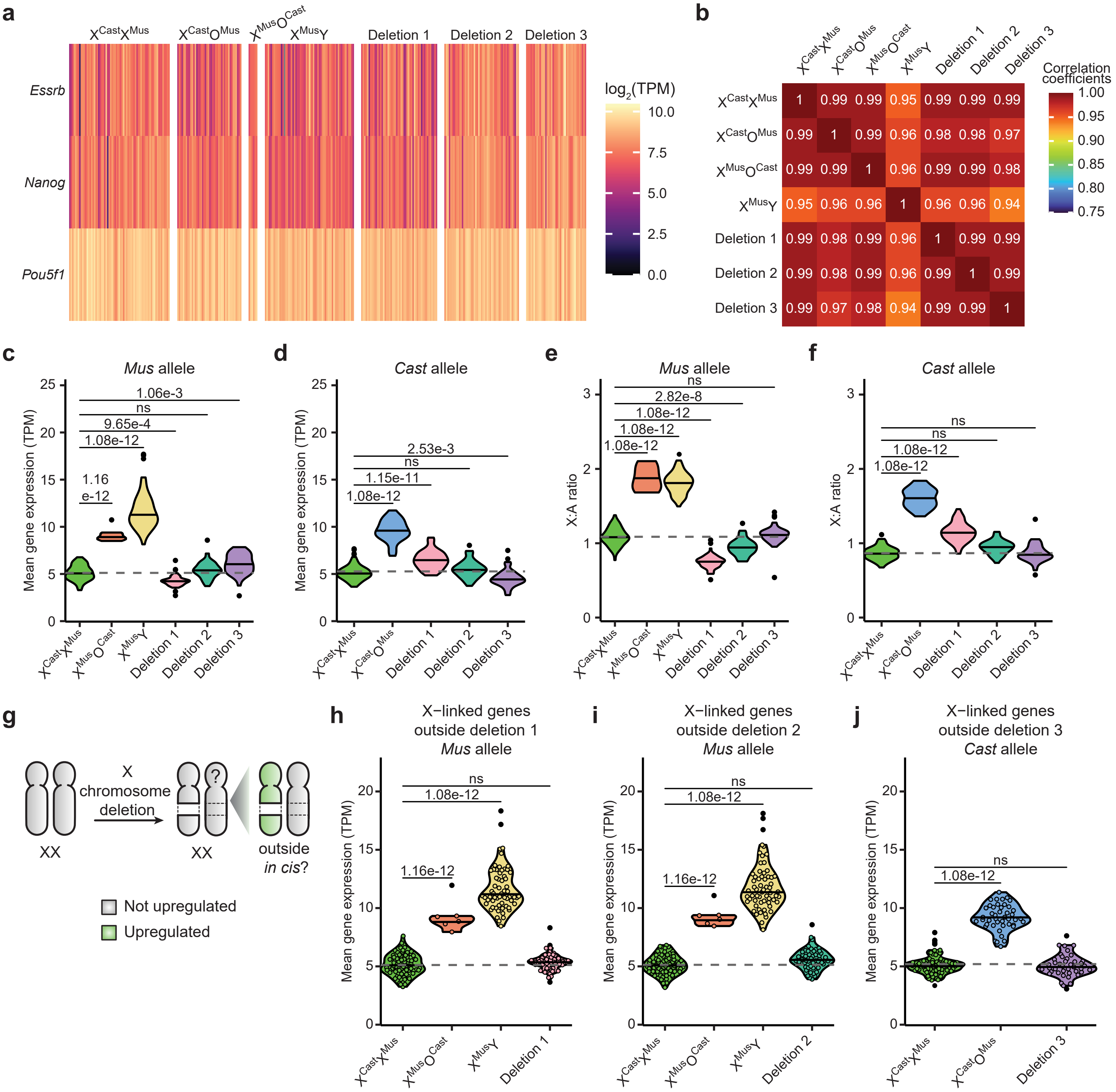

**Supplementary Fig. 5. Genomic and transcriptomic characterisation of cell lines with heterozygous deletions from scRNA-seq experiment.**

**a.** Heatmap showing expression of pluripotency genes *Essrb*, *Nanog* and *Pou5f1*.

**b.** Correlation heatmap showing the Pearson correlation coefficients of the mean total allelic expression of autosomal genes between all cell lines used in the scRNA-seq experiment.

**c, d.** Violin plots separated by cell line show the mean gene expression from the *Mus* (**c**) and *Cast* (**d**) allele of expressed X-linked genes. Dashed line indicates the mean value of XX cells. Each dot represents a cell.

**e, f.** Violin plots separated by cell line show the mean X-to-Autosome ratio (X:A ratio) from the *Mus* (**e**) and *Cast* (**f**) allele of expressed X-linked genes. Dashed line indicates the mean value of XX cells. Each dot represents a cell.

**g**. Schematic depiction of the “XCU center” hypothesis, such that deletion of a region of one X chromosome induces transcriptional upregulation for genes located outside of the deletion region *in cis*.

**h-j.** Violin plots separated by cell line show the mean gene expression from the *Mus* (**h, i**) and *Cast* (**j**) allele of genes located outside of Deletion 1 (**h**), Deletion 2 (**i**), and Deletion 3 (**j**). The dashed line indicates the mean value of $X^{Cast}X^{Mus}$ cells, serving as a reference line for expected expression without upregulation. Each dot represents a cell.

P-values were calculated via the Tukey HSD test (ns - not significant).

### Supplementary Figure 6

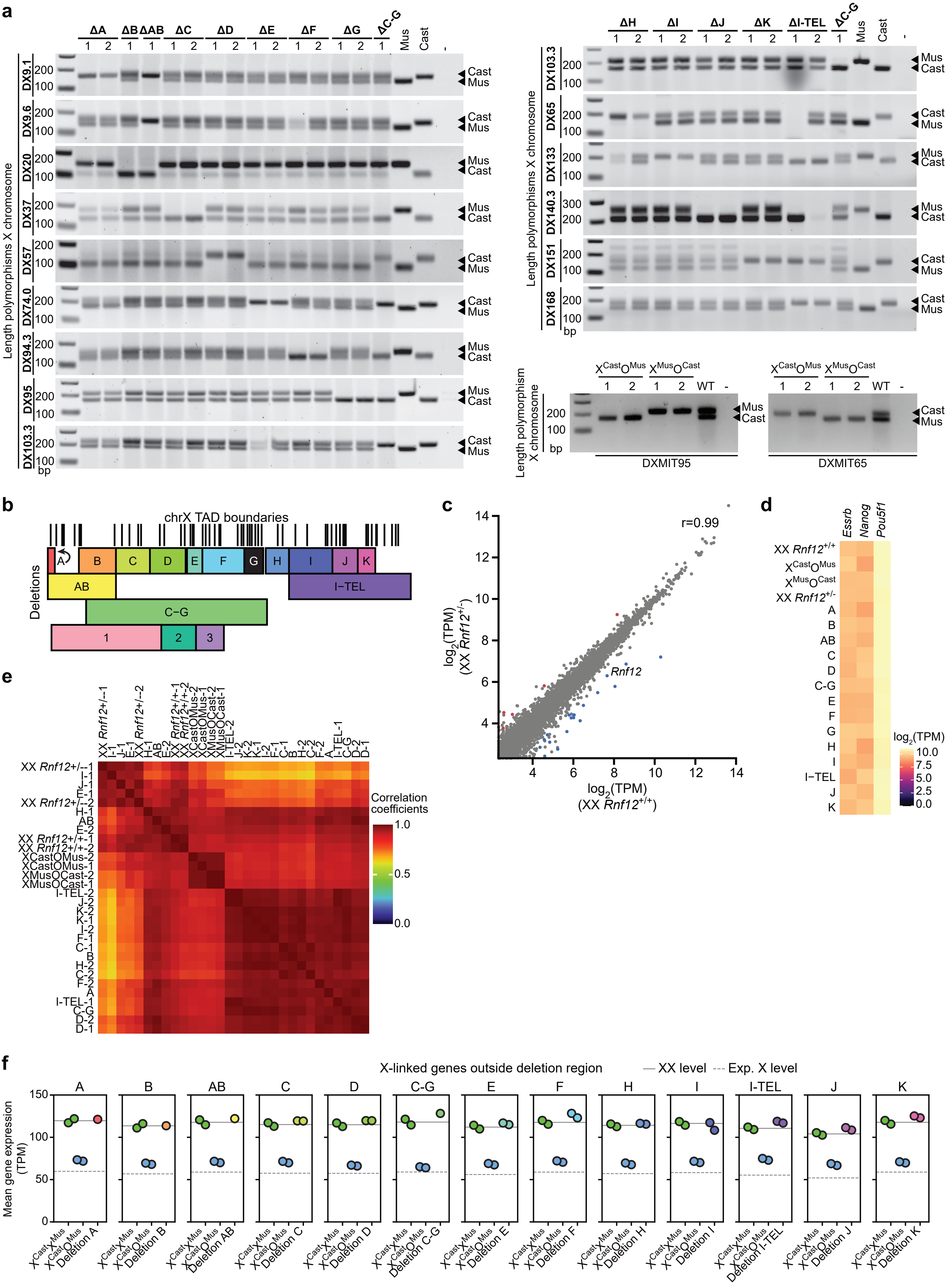

**Supplementary Fig. 6. Genomic and transcriptomic characterization of cell lines with heterozygous deletions from bulk RNA-seq experiment.**

**a.** Genotyping gels validating the deletions in the Deletion A, B, AB, C, D, C-G, E, F, G, H, I, I-TEL, J, K cell lines as well as $X^{Cast}O^{Mus}$ and $X^{Mus}O^{Cast}$ cell lines used for the bulk RNA-seq experiment.

**b.** Schematic depiction of the deleted regions relative to TAD boundary positions (data from Dixon et al., 2012[60]).

**c.** Scatter plot comparing gene expression in transcripts per million (TPM) between WT XX and *Rnf12*$^{+/-}$ XX samples. Each dot represents a gene. Significantly down- and upregulated genes are highlighted in blue and red, respectively. The Pearson correlation coefficient between WT and *Rnf12*$^{+/-}$ XX is displayed.

**d.** Heatmap of log2 TPM values of pluripotency genes *Essrb*, *Nanog* and *Pou5f1* for each sample in the bulk RNA-seq experiment.

**e.** Correlation heatmap showing the Pearson correlation coefficients of the expression of autosomal genes between all cell lines used in the bulk RNA-seq experiment.

**f.** Scatter plots separated by deletion show the mean expression levels of genes located outside each deletion for $X^{Cast}X^{Mus}$, $X^{Cast}O^{Mus}$, and cells with the corresponding deletion. The solid line represents the average expression level from the XX samples, while the dashed line indicates 50% of the $X^{Cast}X^{Mus}$ expression level, serving as a reference line for expected expression without upregulation. Each dot represents a sample

### Supplementary Figure 7

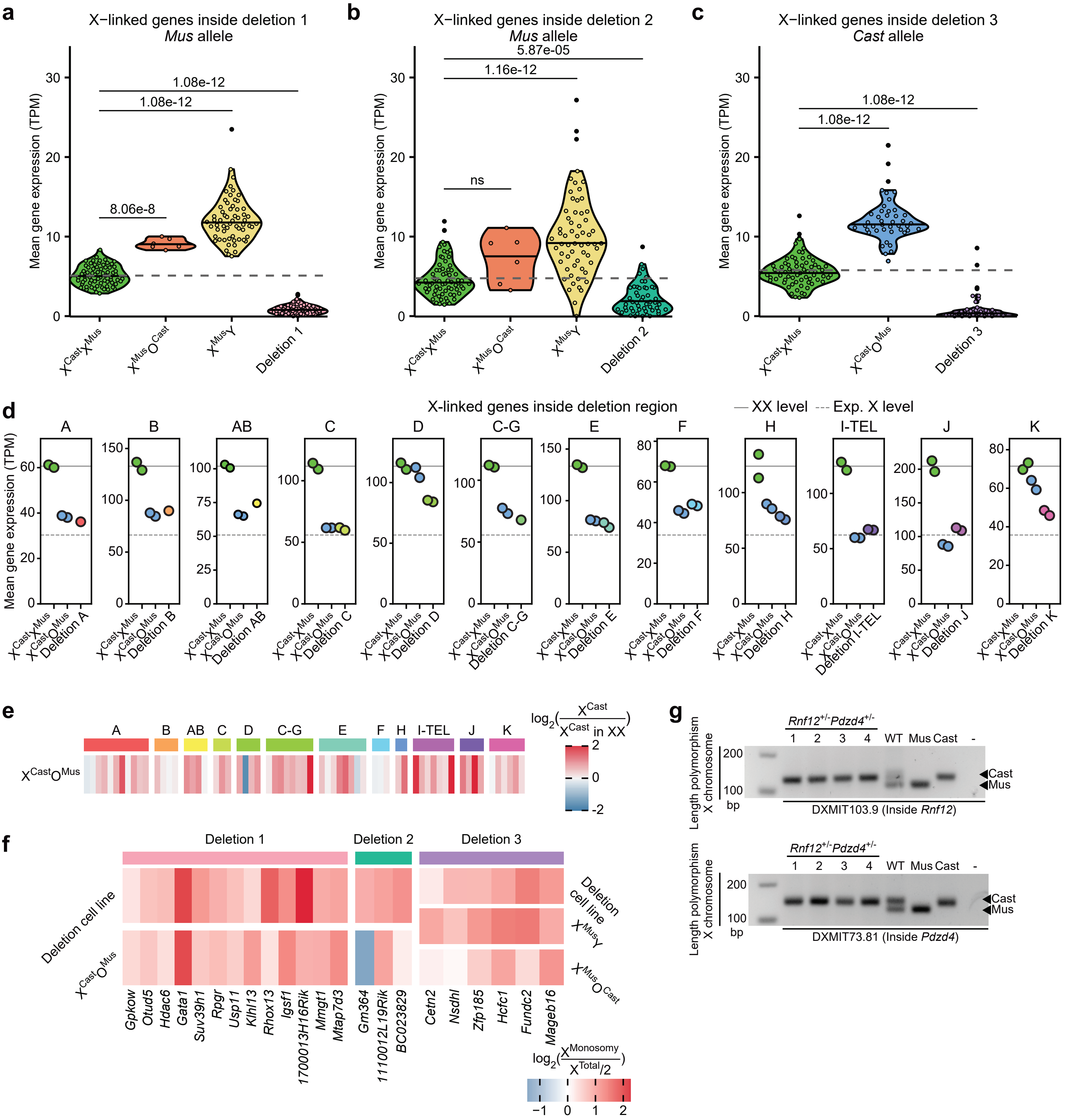

**Supplementary Fig. 7. Transcriptomic characterization of cell lines with heterozygous deletions using genes located inside the deleted regions.**

**a-c.** Violin plots separated by cell line show the mean gene expression from the non-deleted allele inside the region of Deletion 1 (a, *Mus* allele), Deletion 2 (b, *Mus* allele), and Deletion 3 (c, *Cast* allele). The dashed line indicates the mean value of XX cells, serving as a reference line for expected expression without upregulation. Each dot represents a cell. P-values were calculated via the Tukey HSD test (ns - not significant).

**d.** Scatter plots separated by deletion show the mean expression levels of genes located within each deletion for $X^{Cast}X^{Mus}$, $X^{Cast}O^{Mus}$, and cells with the corresponding deletion. The solid line represents the average expression level from the XX samples, while the dashed line indicates 50% of the $X^{Cast}X^{Mus}$ expression level, serving as a reference line for expected expression without upregulation. Deletion I did not contain sufficient expressed genes and has been omitted. Each dot represents a sample.

**e.** Heatmap showing the fold change in $X^{Cast}$ expression in $X^{Cast}O^{Mus}$. Only upregulated genes identified from the non-allelic analysis between $X^{Cast}O^{Mus}$ and $X^{Cast}X^{Mus}$ are shown. Genes are ordered by genomic location and labeled according to the deletion region.

**f.** Heatmap showing the fold change in expression of genes within each deletion and significantly upregulated in cells with the deletion compared to $X^{Cast}X^{Mus}$. Fold change is calculated between Cast expression from either $X^{Cast}O^{Mus}$, Deletion 1, or Deletion 2 cells and total allelic expression divided by 2 from XX cells, or Mus expression from $X^{Mus}O^{Cast}$, $X^{Mus}Y$, or Deletion 3 cells and total allelic expression divided by 2 from $X^{Cast}X^{Mus}$ cells.

**g.** Genotyping gels validating the heterozygous deletion of *Pdzd4* in the *Rnf12*$^{+/-}$ parental line.

### Supplementary Figure 8

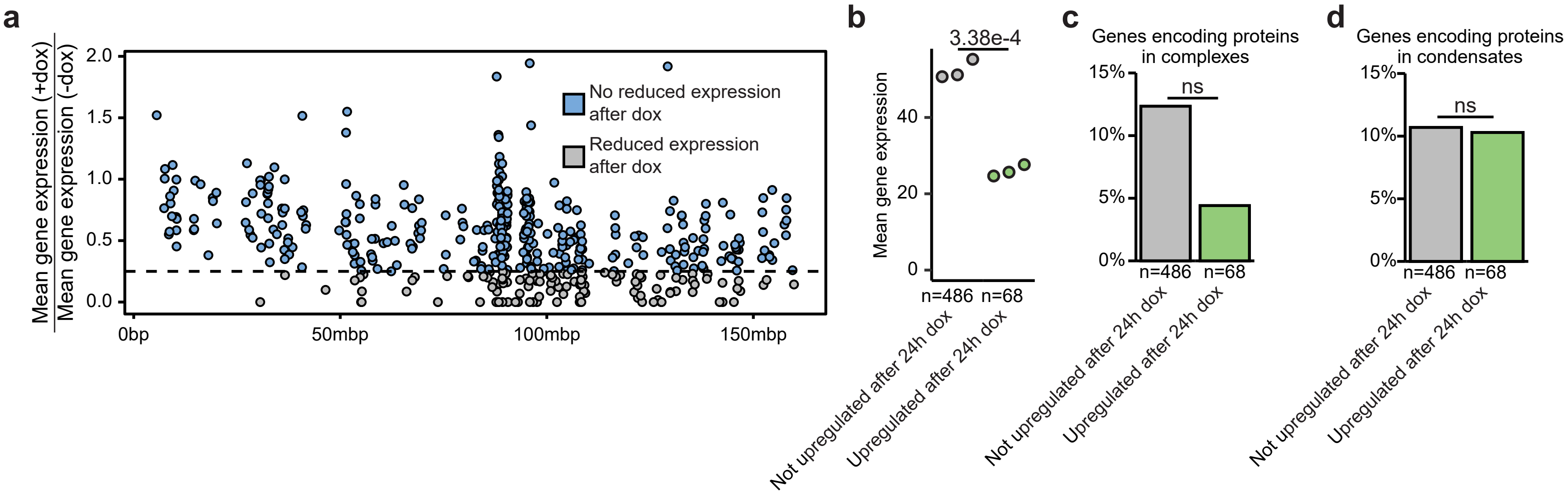

**Supplementary Fig. 8. Characterization of chromosome 3 genes with reduced expression.**

**a.** Scatter plot of the fold change of expression of chromosome 3 genes before and after addition of doxycycline for 24 hours. The dashed line indicates a fold change of 0.25. All genes which fall below the dashed line are defined as genes with reduced expression. Each dot represents a gene.

**b**. Normalized expression of genes annotated as upregulated (n=68, green) or non-upregulated (n=486, gray) identified on chromosome 3 in $X^{Cast}X^{Mus}$ cells prior to addition of doxycycline (ns - not significant). Each dot represents a sample.

**c, d**. Bar plots showing the proportion of genes which code for complex-forming proteins (**c**) or proteins which form condensates (**d**) in the upregulated (n=68, green) and non-upregulated (n=486, gray) gene lists identified on chromosome 3. Each dot represents a sample.

### Supplementary Figure 9

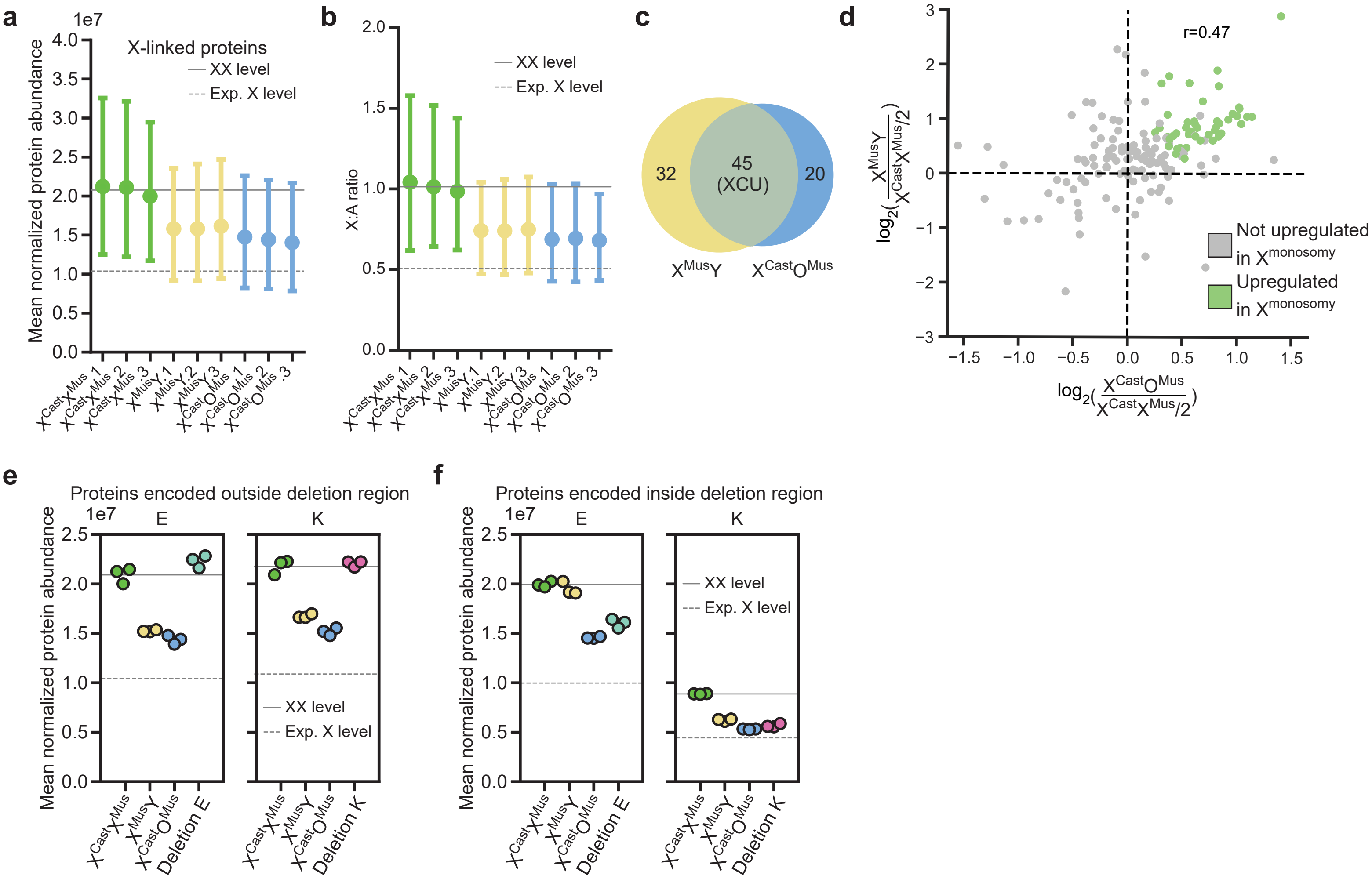

**Supplementary Fig. 9. Characterization of proteomics samples.**

**a, b.** Mean normalized protein abundance (**a**) and X:A ratio (**b**) for all X-linked proteins in XX, XY and $X^{Cast}O^{Mus}$. The mean of each sample is depicted with a dot, with the corresponding 95% confidence intervals shown as error bars. The solid line represents the average abundance levels from the XX samples, while the dashed line indicates 50% of the XX level, serving as a reference line for expected abundance without upregulation.

**c.** Venn diagram showing overlap between proteins upregulated in $X^{Mus}Y$ (yellow) and $X^{Cast}O^{Mus}$ (blue) samples. The 45 overlapping proteins were used in subsequent analyses of upregulated proteins.

**d.** Scatter plot of the $log_2$ fold change in copy number-corrected protein abundance in both $X^{Cast}O^{Mus}$ and $X^{Mus}Y$ samples compared to controls. Proteins are colored by upregulation status, and Pearson correlation coefficient is shown. Each dot represents a protein.

**e, f.** Scatter plots separated by region showing the mean normalized abundance levels of proteins encoded by genes located outside (**e**) and within (**f**) region E and K for XX, XY, $X^{Cast}O^{Mus}$ and cells with the corresponding deletion. The solid line represents the average abundance levels from the XX samples, while the dashed line indicates 50% of the XX level, serving as a reference line for expected abundance without upregulation. Each dot represents a sample.

**Supplementary Table 1. Upregulated genes**

| Gene name | Ensembl ID |
|---|---|
| Tfe3 | ENSMUSG00000000134 |
| Dlg3 | ENSMUSG00000000881 |
| Uba1 | ENSMUSG00000001924 |
| Pdzd4 | ENSMUSG00000002006 |
| Pola1 | ENSMUSG00000006678 |
| Pls3 | ENSMUSG00000016382 |
| Fhl1 | ENSMUSG00000023092 |
| Maoa | ENSMUSG00000025037 |
| Fundc1 | ENSMUSG00000025040 |
| Nr0b1 | ENSMUSG00000025056 |
| Tbl1x | ENSMUSG00000025246 |
| Hsd17b10 | ENSMUSG00000025260 |
| Gnl3l | ENSMUSG00000025266 |
| Pfkfb1 | ENSMUSG00000025271 |
| Tro | ENSMUSG00000025272 |
| Sat1 | ENSMUSG00000025283 |
| Prdx4 | ENSMUSG00000025289 |
| Kdm5c | ENSMUSG00000025332 |
| Rps6ka6 | ENSMUSG00000025665 |
| Tmem47 | ENSMUSG00000025666 |
| Ndufb11 | ENSMUSG00000031059 |
| Gpc4 | ENSMUSG00000031119 |
| Plp2 | ENSMUSG00000031146 |
| Otud5 | ENSMUSG00000031154 |
| Pim2 | ENSMUSG00000031155 |
| Eras | ENSMUSG00000031160 |
| Gata1 | ENSMUSG00000031162 |
| Glod5 | ENSMUSG00000031163 |
| Rbm3 | ENSMUSG00000031167 |
| Porcn | ENSMUSG00000031169 |
| Fundc2 | ENSMUSG00000031198 |
| Mtcp1 | ENSMUSG00000031200 |
| Msn | ENSMUSG00000031207 |
| Awat2 | ENSMUSG00000031220 |
| Itm2a | ENSMUSG00000031239 |
| Chmp1b2 | ENSMUSG00000031242 |
| Hmgn5 | ENSMUSG00000031245 |
| Acsl4 | ENSMUSG00000031278 |
| Slc7a3 | ENSMUSG00000031297 |
| Pdha1 | ENSMUSG00000031299 |
| Nlgn3 | ENSMUSG00000031302 |
| Nono | ENSMUSG00000031311 |
| Itgb1bp2 | ENSMUSG00000031312 |
| Rps4x | ENSMUSG00000031320 |
| Flna | ENSMUSG00000031328 |
| Gabre | ENSMUSG00000031340 |
| Cetn2 | ENSMUSG00000031347 |
| Syap1 | ENSMUSG00000031357 |
| Msl3 | ENSMUSG00000031358 |
| Hcfc1 | ENSMUSG00000031386 |
| Naa10 | ENSMUSG00000031388 |
| Plxna3 | ENSMUSG00000031398 |
| G6pdx | ENSMUSG00000031400 |
| Dkc1 | ENSMUSG00000031403 |
| Slc10a3 | ENSMUSG00000032806 |
| Armcx2 | ENSMUSG00000033436 |
| Armcx1 | ENSMUSG00000033460 |
| Fndc3c1 | ENSMUSG00000033737 |
| Slc16a2 | ENSMUSG00000033965 |
| Ogt | ENSMUSG00000034160 |
| Kif4 | ENSMUSG00000034311 |
| Eda2r | ENSMUSG00000034457 |
| Pdk3 | ENSMUSG00000035232 |
| Pcyt1b | ENSMUSG00000035246 |
| Tab3 | ENSMUSG00000035476 |
| Fthl17a | ENSMUSG00000035491 |
| Pabir2 | ENSMUSG00000036022 |
| Zfp280c | ENSMUSG00000036916 |
| Kdm6a | ENSMUSG00000037369 |
| Suv39h1 | ENSMUSG00000039231 |
| Bcor | ENSMUSG00000040363 |
| Gemin8 | ENSMUSG00000040621 |
| Tspyl2 | ENSMUSG00000041096 |
| Smc1a | ENSMUSG00000041133 |
| Ptchd1 | ENSMUSG00000041552 |
| Kctd12b | ENSMUSG00000041633 |
| Rragb | ENSMUSG00000041658 |
| Amot | ENSMUSG00000041688 |
| Alg13 | ENSMUSG00000041718 |
| Pwwp3b | ENSMUSG00000042515 |
| Fam199x | ENSMUSG00000042595 |
| Foxo4 | ENSMUSG00000042903 |
| Slc25a53 | ENSMUSG00000044348 |
| Tceal3 | ENSMUSG00000044550 |
| Shroom2 | ENSMUSG00000045180 |
| Hnrnph2 | ENSMUSG00000045427 |
| Bex3 | ENSMUSG00000046432 |
| Nexmif | ENSMUSG00000046449 |
| Spin2c | ENSMUSG00000046550 |
| Tmem164 | ENSMUSG00000047045 |
| Arxes2 | ENSMUSG00000048040 |
| Cypt3 | ENSMUSG00000048573 |
| Rtl5 | ENSMUSG00000049191 |
| Lpar4 | ENSMUSG00000049929 |
| Ubqln2 | ENSMUSG00000050148 |
| Ercc6l | ENSMUSG00000051220 |
| Nrk | ENSMUSG00000052854 |
| Rlim | ENSMUSG00000056537 |
| Tspan7 | ENSMUSG00000058254 |
| Nhs | ENSMUSG00000059493 |
| Mmgt1 | ENSMUSG00000061273 |
| Hs6st2 | ENSMUSG00000062184 |
| Brwd3 | ENSMUSG00000063663 |
| Utp14a | ENSMUSG00000063785 |
| Med14 | ENSMUSG00000064127 |
| Zic3 | ENSMUSG00000067860 |
| Map7d3 | ENSMUSG00000067878 |
| Shroom4 | ENSMUSG00000068270 |
| Ccdc160 | ENSMUSG00000073207 |
| Nudt11 | ENSMUSG00000073295 |
| Rab9 | ENSMUSG00000079316 |
| Rpl36a | ENSMUSG00000079435 |
| Tmsb15b2 | ENSMUSG00000089996 |

**Supplementary Table 2. Metadata of cell lines**

| Deletion | ChrX coordinates Start | ChrX coordinates End | Size (Mbp) | Number of genes in deletion Total | Number of genes in deletion Expressed | Number of genes in deletion Upregulated | % upregulated |
|---|---|---|---|---|---|---|---|
| 1 | 7,640,796 | 57,081,805 | 49.4 | 308 | 120 | 13 | 10.8 |
| 2 | 57,075,843 | 72,687,204 | 15.6 | 52 | 27 | 3 | 11.1 |
| 3 | 72,683,014 | 85,235,510 | 12.6 | 102 | 62 | 7 | 11.3 |
| A | 5,977,301 | 9,126,867 | 3.1 | 65 | 33 | 7 | 21.2 |
| B | 19,965,047 | 36,657,607 | 16.7 | 80 | 21 | 4 | 19.0 |
| AB | 5,977,301 | 36,657,607 | 30.7 | 187 | 69 | 18 | 26.1 |
| C | 36,657,607 | 51,952,154 | 15.3 | 99 | 39 | 8 | 20.5 |
| D | 51,952,154 | 68,014,381 | 16.1 | 64 | 22 | 4 | 18.2 |
| C-G | 23,219,171 | 104,671,164 | 81.5 | 461 | 176 | 38 | 21.6 |
| E | 68,651,334 | 75,601,939 | 7.0 | 103 | 52 | 13 | 25.0 |
| F | 75,567,564 | 94,452,400 | 18.9 | 58 | 16 | 2 | 12.5 |
| G | 94,460,524 | 102,939,806 | 8.5 | 69 | 34 | NA | NA |
| H | 104,088,048 | 114,560,591 | 10.5 | 48 | 19 | 3 | 15.8 |
| I | 114,561,262 | 134,111,786 | 19.6 | 36 | 4 | 2 | 50.0 |
| J | 134,112,553 | 145,486,844 | 11.4 | 105 | 45 | 11 | 24.4 |
| K | 145,506,466 | 153,393,474 | 7.9 | 44 | 26 | 8 | 30.8 |
| I-TEL | 114,561,262 | 169,370,103 | 54.8 | 275 | 115 | 33 | 28.7 |

**Supplementary Table 3. Upregulated proteins**

| Protein name | Ensembl ID |
|---|---|
| UBE2A | ENSMUSG00000016308 |
| PLS3 | ENSMUSG00000016382 |
| CUL4B | ENSMUSG00000031095 |
| MED14 | ENSMUSG00000064127 |
| MED12 | ENSMUSG00000079487 |
| SHROOM2 | ENSMUSG00000045180 |
| LAS1L | ENSMUSG00000057421 |
| FLNA | ENSMUSG00000031328 |
| RPL10 | ENSMUSG00000008682 |
| KDM6A | ENSMUSG00000037369 |
| RBM3 | ENSMUSG00000031167 |
| UBL4A | ENSMUSG00000015290 |
| MSN | ENSMUSG00000031207 |
| PDHA1 | ENSMUSG00000031299 |
| GDI1 | ENSMUSG00000015291 |
| VBP1 | ENSMUSG00000031197 |
| RPS4X | ENSMUSG00000031320 |
| RPL39 | ENSMUSG00000079641 |
| IDH3G | ENSMUSG00000002010 |
| SMS | ENSMUSG00000071708 |
| CDK16 | ENSMUSG00000031065 |
| GSPT2 | ENSMUSG00000071723 |
| RBBP7 | ENSMUSG00000031353 |
| HCFC1 | ENSMUSG00000031386 |
| DDX3X | ENSMUSG00000000787 |
| ZIC3 | ENSMUSG00000067860 |
| UTP14A | ENSMUSG00000063785 |
| SMARCA1 | ENSMUSG00000031099 |
| GNL3L | ENSMUSG00000025266 |
| CSTF2 | ENSMUSG00000031256 |
| RAP2C | ENSMUSG00000050029 |
| PQBP1 | ENSMUSG00000031157 |
| NONO | ENSMUSG00000031311 |
| SMC1A | ENSMUSG00000041133 |
| PDZD11 | ENSMUSG00000015668 |
| PHF6 | ENSMUSG00000025626 |
| MCTS1 | ENSMUSG00000000355 |
| APOO | ENSMUSG00000079508 |
| DKC1 | ENSMUSG00000031403 |
| CCDC22 | ENSMUSG00000031143 |
| ACSL4 | ENSMUSG00000031278 |
| MORF4L2 | ENSMUSG00000031422 |
| RBMX | ENSMUSG00000031134 |
| HDAC6 | ENSMUSG00000031161 |

**Supplementary Table 4. List of gRNA sequences**

| Name | Sequence (5' - 3') | Additional info | Reference |
|---|---|---|---|
| gRNA_Del_A-5' | TTAAGCACATACTACCATAG | | |
| gRNA_Del_A-3' | TCTGGCTTCTTTCAATACAG | | |
| gRNA_Del_B-5' | TGACAAGTGCTAATGAACAA | | |
| gRNA_Del_B-3'_Del_C-5' | TTTCTGGGTATGATTTACCA | | |
| gRNA_Del_C-3'_Del_D-5' | CAGGGGCATGGACTGCCGAG | | |
| gRNA_Del_D-3' | CCATTGCTAATGTGTCCATG | | |
| gRNA_Del_E-5' | TTAATAATATGCTAAACCAG | | |
| gRNA_Del_E-3' | TTATGGTTGCAGTTTGCGGG | | |
| gRNA_Del_F-5' | AATCACCTGGTATGAGACGG | | |
| gRNA_Del_F-3' | GGAGACCACCATGACCGGGT | | |
| gRNA_Del_G_5' | GCTTAGTGTTCAGACAACGG | | |
| gRNA_Del_G_3' | CAGGCATGAGGTTTCCGTAG | | |
| gRNA_Del_H-5' | GCACTATTAGCAGAATGATC | | |
| gRNA_Del_H-3' | ATATTGTTTTGTCATTAGGG | | |
| gRNA_Del_I-5' | TAGTGCTATTTACATTAGCA | | |
| gRNA_Del_I-3' | GTCAGAAGTTTCTCTCCCGA | | |
| gRNA_Del_J-5' | CAGAATGCTTCCAGACGGAT | | |
| gRNA_Del_J-3' | GAGCATCGCCTGAACCAGCC | | |
| gRNA_Del_K-5' | CAGTTGGGGGGATACGGTTT | | |
| gRNA_Del_K-3' | GAGTCTCATGCTAACCGTGA | | |
| gRNA_Rnf12_5'-1 | AAAGCGCTGTACAAAAAGTT | Rnf12+/- line for del A to I & K, Rnf12+/-Pdzd4+ | 1 |
| gRNA_Rnf12_3'-1 | GGAACAAGTACTCTAAACTA | | 1 |
| gRNA_Rnf12_5'-2 | CATATTGAACTGTACTAAAG | Rnf12+/- line del J and I-TEL | |
| gRNA_Rnf12_3'-2 | TTACTTGGAAACTACAATGC | | |
| gRNA_Pdzd4_5'-1 | ATGCTAGGGCCCAAGCAATG | 5'-1 and 3'-2 gRNAs for clone 1 and 2 | |
| gRNA_Pdzd4_3'-2 | ATTGGCATTATCTGGTCTCA | | |
| gRNA_Pdzd4_5'-2 | CAACAACCCACCCCATTGCT | 5'-2 and 3'-1 gRNAs for clone 3 and 4 | |
| gRNA_Pdzd4_3'-1 | GGGCCTCAGGAGTCGGCGCA | | |

**Supplementary Table 5. List of genotyping primers**

| Name | Sequence (5' - 3') | Description | Reference |
|---|---|---|---|
| DXMIT9.1_Fw | TCCTTTTCCCACTGACAGGC | Inside deletion A | |
| DXMIT9.1_Rv | CAAAGCCTCCACGACTTCCT | | |
| DXMIT9.6_Fw | CCTGTGTAAGCAAGTGCAGC | Inside deletion AB | |
| DXMIT9.6_Rv | CAGCTGCCATCCCTTCCTTT | | |
| DXMIT20.0_Fw | CCAGCAATCCTTTGTCACTTCC | Inside deletion B | |
| DXMIT20.0_Rv | GCAGGCTTGTGCATGCTAGT | | |
| DXMIT37_Fw | TCCCAGACATTCGACCAAACA | Inside deletion C | |
| DXMIT37_Rv | TACAAATGGCTGGTCCTCCA | | |
| DXMIT57_Fw | ACCCTAGCCGGTTACTATCTCC | Inside deletion D | |
| DXMIT57_Rv | TCTCTCTCATTCCCCCCC | | |
| DXMIT74.0_Fw | AAGCACAAAGATCGGTCAGG | Inside deletion E | |
| DXMIT74.0_Rv | CGCAACACACATGTACACACA | | |
| DXMIT94.3_Fw | CTATGACCAGCAAGAGTGGCT | Inside deletion F | |
| DXMIT94.3_Rv | ACTTCCCTCCAGGGTAGCTT | | |
| DXMIT95_Fw | CCCTGAGGCTGGGAGTCTAT | Inside deletion G | |
| DXMIT95_Rv | AGGACGTAAGGAGAGTTAGAGAGA | | |
| DXMIT103.3_Fw | AGCACTCATGGAGCCCAAAA | Inbetween deletion G and H | |
| DXMIT103.3_Rv | AGCCAACACAGTGCCCATAA | | |
| DXMIT65_Fw | ATATTAAGGGAGGTAACAAAGACCC | Inside deletion H | Whitehead institute at MIT; Center for genomic research |
| DXMIT65_Rv | GGTTTCTGTGATTGCTATAGGACA | | |
| DXMIT133_Fw | TGGGGCTCAGTGGAAAAACA | Inside deletion I | |
| DXMIT133_Rv | GCTGGTAGTGACAGATCACTTG | | |
| DXMIT140.3_Fw | ACAGGACACCACGACTTAGT | Inside deletion J | |
| DXMIT140.3_Rv | GTGGGAGAGGAAGCAGAGAC | | |
| DXMIT151_Fw | TGTTCTATATTGCTTTGTTAGGTTTT[illegible] | Inside deletion K | |
| DXMIT151_Rv | GCAAAAAAGAAACCAACCCA | | |
| DXMIT168_Fw | GTCTGGATTTGGGTTCTAAATATTT | Outside deletion K 3' | |
| DXMIT168_Rv | AAGCACATACACACATACGTTCTC | | |
| DXMIT103.9_Fw | GCCAGCCTGGTCTACAAAGT | Inside Rnf12 | |
| DXMIT103.9_Rv | AGGAGATGCAAAATGGCAGT | | |
| DXMIT73.81_Fw | CAGGACACATGCATCTTGGG | Inside Pdzd4 | |
| DXMIT73.81_Rv | GGAAAGTGAGCAAGACTGGCA | | |

**Supplementary Table 6. List of allele-specific qPCR primers**

| Name | Sequence (5' - 3') | Description | Reference |
|---|---|---|---|
| Rplp0_qPCR_FW | TCCAGAGGCACCATTGAAATT | | 2 |
| Rplp0_qPCR_RV | TCGCTGGCTCCCACCTT | | 2 |
| H2afz_qPCR_FW | GGCCGTATTCATCGACACCTGA | | Origene, MP205904 |
| H2afz_qPCR_RV | GACGCATTTCCTGCCAACTCAAG | | Origene, MP205904 |
| PDZD4_129_F1 | TCCCTTCTTGGGGCTTTCTATT | SNP rs31667483 PDZD4 3'UTR | |
| PDZD4_Cast_F1 | TCCCTTCTTGGGGCTTTCTATC | SNP rs31667483 PDZD4 3'UTR | |
| PDZD4_R1 | ACCCTCACCCTCCCTTAGAA | SNP rs31667483 PDZD4 3'UTR | |
